# Self-blinking dye restores efficient use of nanobodies in single-molecule localization microscopy

**DOI:** 10.64898/2025.12.17.694880

**Authors:** Samrat Basak, Kaushik Inamdar, Yoav G. Pollack, László Albert, Daniel C. Jans, Stefan Jakobs, Jörg Enderlein, Roman Tsukanov, Felipe Opazo

## Abstract

Direct stochastic optical reconstruction microscopy (dSTORM) relies on controlled fluorophore blinking to achieve nanometer-scale resolution, yet the field’s benchmark dye, Alexa Fluor 647, underperforms when conjugated to nanobodies, limiting the practical use of minimal-linkage labeling strategies. Here, we show that the self-blinking dye JF635b overcomes this limitation by maintaining robust, photostable blinking upon conjugation to nanobodies under buffer-independent conditions. This enables reliable single-molecule localization microscopy without the need for complex switching buffers. Using JF635b-labeled nanobodies, we demonstrate consistent performance across multiple imaging modalities, including wide-field dSTORM, fluorescence lifetime SMLM, and MINFLUX nanoscopy, achieving localization precisions from ∼15 nm down to the sub-nanometer regime. In addition, JF635b supports long-term sample preservation and efficient blinking even in pure water, enabling minimally perturbative imaging conditions. Together, these results establish self-blinking dSTORM as a robust and accessible platform for quantitative nanoscopy across experimental contexts.

## Introduction

Single-molecule localization microscopy (SMLM)^1^ enables nanometer-scale localization precision. Among its core implementations, direct stochastic optical reconstruction microscopy (dSTORM)^2^ routinely achieves localization precision below 20 nm, but performance critically depends on the fluorophore blinking characteristics. Alexa Fluor 647 (AF647) has long been the benchmark dye, exhibiting robust blinking in buffers such as GLOX-ME^2^. The reductive properties of the blinking buffers alter disulfide bonds in antibodies and nanobodies, weakening them or rendering them dysfunctional. Therefore, careful optimization of buffer composition for each sample is required, especially in multitarget imaging, which often requires compromises, leading to the development of sequential-labeling strategies such as NanoPlex^3^ or microfluidics-enhanced Exchange-PAINT^4^.

Despite its widespread adoption, dSTORM remains highly sensitive to imaging conditions, with fluorophore blinking typically requiring carefully optimized chemical environments. Small variations in buffer composition, oxygen-scavenging efficiency, or reducing agents can substantially affect blinking behavior, localization density, and photostability, posing a major challenge to reproducibility across laboratories and imaging facilities. These constraints are particularly problematic for multitarget imaging, long acquisition times, and workflows involving repeated imaging or sample storage.

Linkage error in indirect immunofluorescence (IF) labeling occurs when the primary antibody (1.Ab) targeting the protein of interest (POI) is labeled with a fluorescent secondary antibody (2.Ab). Indirect IF displaces fluorophores by up to 25 nm from the POI. Nanobodies (Nbs) reduce this linkage error to approximately 3 nm, making them ideal probes^5^ for SMLM nanoscopy. However, a practical limitation has emerged: when directly conjugated to nanobodies, commonly used dyes such as Alexa Fluor 647 often exhibit compromised blinking behavior and reduced photostability. This undermines the very advantages nanobodies offer for high-precision SMLM and restricts their use in workflows that rely on consistent and sustained photoswitching.

Self-blinking dyes provide an elegant solution by spontaneously cycling between bright and dark states without the need for blinking-reducing buffers. Several such fluorophores have been explored for super-resolution microscopy, including specialized dyes for live-cell super-resolution imaging^6^, Thioflavin T for fibrils^7^, rhodamine-coumarin hybrids^8^, silicon rhodamine analogs for STORM-MINFLUX^9^, and rhodamine-based dyes for SOFI^10^.

JF635b is a newly developed, spontaneously blinking rhodamine from the hydroxymethyl-SiR Janelia Fluor series, described in Holland et al.^11^. It exhibits an excellent duty cycle, is ideally suited to SMLM applications, and operates without reducing-blinking buffers, significantly simplifying the imaging workflow and making it compatible with a wide range of labeling strategies. Although its photophysical properties have been characterized when conjugated to HaloTags^11^, its versatility and potential for SMLM for optimal blinking and minimal linkage error due to conjugation to different types of Nbs have not been explored. Here, we benchmark JF635b against AF647 across Ab and Nb conjugates using wide-field self-blinking dSTORM (sb-dSTORM), confocal FL-SMLM^12,13^, and MINFLUX^14^.

## Results

To define how fluorophore choice impacts the performance of nanobody-based single-molecule localization microscopy (SMLM), we benchmarked the blinking behavior of Alexa Fluor 647 (AF647) and the self-blinking dye JF635b in a direct dSTORM implementation. COS-7 cells expressing TOMM70–EGFP–ALFA were immunostained using two widely applied nanobodies, NbGFP (binding EGFP) and NbALFA (binding the ALFA-tag), each conjugated to either AF647 or JF635b. Samples labeled with AF647 were imaged under standard GLOX–MEA switching conditions, whereas JF635b-labeled samples were imaged in phosphate-buffered saline without the use of chemical blinking buffers (Fig. 1).

**Figure 1.**
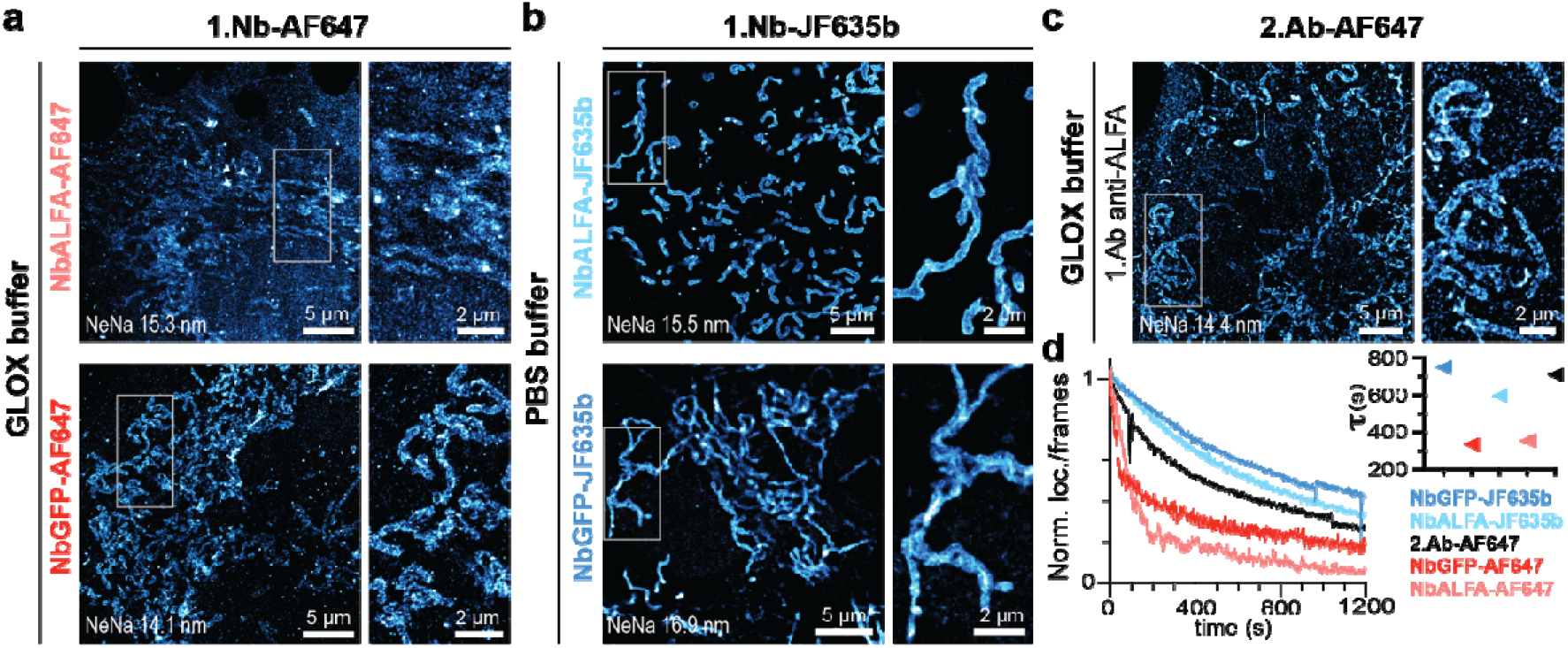
dSTORM using nanobodies conjugated with AF647 or JF635b. Primary nanobody (1.Nb) NbALFA and NbGFP, each conjugated with 2 AF647 (**a**) or JF635b (**b**), were used to directly label transfected COS-7 cells displaying EGFP and ALFA-tag on their mitochondria. (**c**) As a control, conventional 1.Ab anti-ALFA-tag, revealed by secondary antibodies (2.Ab-AF647). Samples with AF647 were imaged under GLOX-MEA buffer, while samples with JF635b were imaged in PBS. Average localization precisions (NeNa) are indicated on the images. (**d**) Localization per frame was plotted, with the blinking differences between conditions shown as tau (τ) for each curve (details provided in Methods).

Nanobodies conjugated to AF647 exhibited strongly impaired blinking behavior, characterized by reduced photon yields, shortened on-times, and rapid photobleaching, with most emitters bleaching within approximately 5 minutes of acquisition in wide-field SMLM under optimal imaging conditions (Supp. Fig. S1 and Supp. Table S1). Compared to a control condition using conventional indirect immunofluorescence to detect the ALFA-tag, NbALFA-AF647 conjugates produced ∼70% fewer localizations, an effect consistently observed for both NbGFP and NbALFA. Although both Nbs display comparable binding affinities, NbGFP-AF647 yielded marginally better reconstructions than NbALFA-AF647, while NbALFA-AF647 showed particularly poor blinking and rapid signal loss, resulting in clearly suboptimal mitochondrial reconstructions (Fig. 1a-b). Importantly, these differences were independent of the target interaction and were consistently observed across nanobody types, indicating that the reduced performance originates from the dye–nanobody conjugate rather than from binding affinity or labeling efficiency.

In contrast, Nbs conjugated to JF635b exhibited stable, sustained self-blinking behavior when imaged in PBS. Individual emission bursts yielded approximately 500–1500 photons with on-times of 40–60 ms and remained detectable over tens of thousands of frames without appreciable photobleaching (Supp. Movies 1–5). Although individual blinking events were shorter-lived than those observed for AF647 under reducing conditions, the markedly higher number of blinking cycles resulted in higher localization densities, enabling faster and more complete reconstructions. As a result, JF635b-labeled Nbs produced continuous, densely sampled super-resolution images with localization precision comparable to or exceeding that obtained with conventional antibody-based labeling, confirming the high photostability and robustness of JF635b under buffer-independent sb-dSTORM conditions (Fig. 1a-d; Supp. Table S1).

To test whether buffer-independent self-blinking nanobody probes are compatible with widely used antibody-based labeling workflows, we next examined secondary nanobody (2.Nb) detection strategies, which offer flexibility, multiplexing capacity, and broad applicability across established primary antibodies^15^. Microtubules in COS-7 cells were labeled using a monoclonal anti-α-tubulin primary antibody, revealed with secondary nanobodies conjugated to two JF635b molecules per nanobody, and imaged by wide-field sb-dSTORM in phosphate-buffered saline (Fig. 2; Supp. Fig. S2 and Supp. Movie 6).

**Figure 2.**
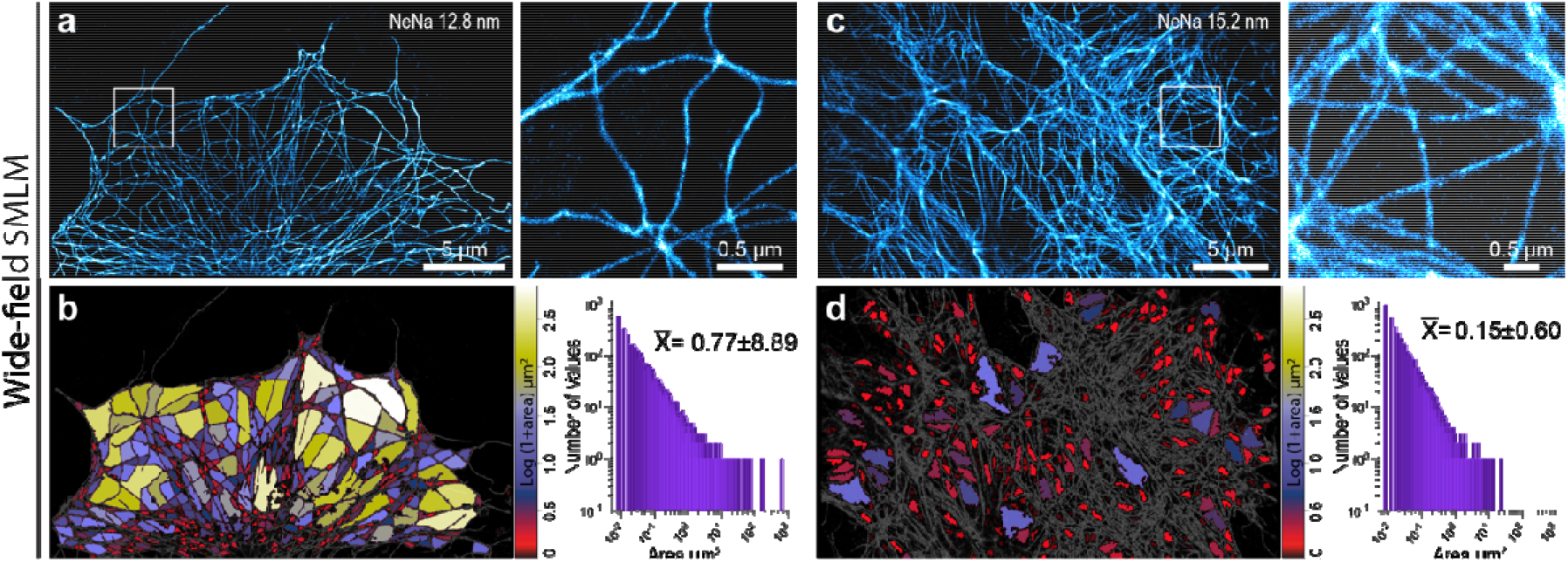
Applications of sb-dSTORM across labeling strategies and SMLM modalities. (**a-d**) Wide-field sb-dSTORM in COS-7 cells labeled with 1.Ab and 2.Nb-JF635b. (**a, c**) Full reconstruction with zoom-ins of microtubule and vimentin. (**b, d**) Segmentation analysis for (a, c) and area histograms for all datasets combined (microtubule, vimentin, respectively).).

JF635b-labeled 2.Nbs yielded continuous and densely sampled reconstructions of microtubule filaments with a mean localization precision of 12.8 ± 4.5 nm (Fig. 2a). The high localization density enabled reliable segmentation of the filament network, revealing a characteristic mesh organization with a mean area of 0.77 ± 8.89 µm^2^ (Fig. 2b; segmentation workflow in Supp. Fig. S3). Applying the same labeling strategy to vimentin intermediate filaments using a polyclonal rabbit primary antibody produced similarly robust reconstructions with a localization precision of 15.2 ± 4.7 nm and, as expected, a denser network architecture characterized by a smaller mean mesh area of 0.15 ± 0.60 µm^2^ (Fig. 2c,d; Supp. Fig. S4 and Supp. Movie 7). Comparable high labeling efficiency and image quality were observed for additional targets, including peroxisomes (Supp. Fig. S5), underscoring the versatility and general applicability of this approach.

### Self-blinking nanobody probes enable robust imaging in a buffer-independent manner

To assess long-term stability, COS-7 samples labeled with a dye were stored in PBS at 4°C and re-imaged after extended periods. Over storage times of up to one year, JF635b-labeled nanobodies maintained stable blinking behavior and localization precision, with only a modest reduction in labeling density observed after 12 months (Fig. 3a). In contrast, AF647 typically exhibits a significant decay in blinking performance within weeks. The pronounced stability of Nb–JF635b conjugates enables long-term sample preservation, repeated reimaging, and high-throughput data acquisition without the need for buffer exchange or relabeling.

**Figure 3.**
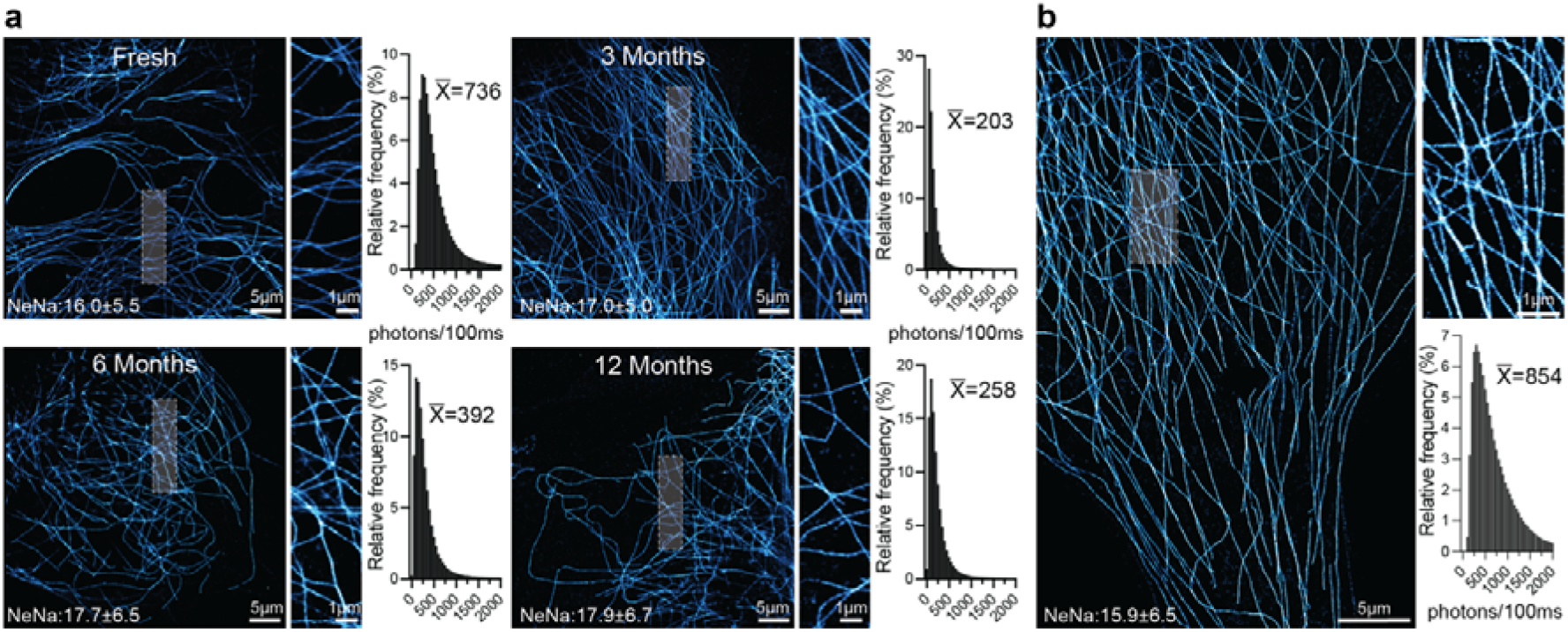
Long-term storage of the sample and the water performance of the Nb-JF635. (**a**) dSTORM images of the same sample imaged immediately after labeling (Fresh) and after 3, 6, and 12 months of storage at +4°C in PBS. Blinking performance and localization precision (NeNa) as well as the photon output of 2.Nb-JF635b remained stable throughout the 12 months. Above the distributions of photons/100 ms, the average (x□) is displayed. (**b**) dSTORM image of microtubules labeled with JF635b and imaged in pure water. The average localization precision (NeNa) and photon output were comparable to or higher than those in PBS.

We next asked whether the intrinsic blinking behavior of JF635b could support SMLM under conditions that are typically incompatible with conventional dSTORM fluorophores. Remarkably, 2.Nb-JF635b conjugates exhibited stable, reversible blinking even when imaged in pure water, in the complete absence of salts or additives (Fig. 3b; Supp. and Supp. Movie 8). Despite these minimally perturbed conditions, localization densities and reconstruction quality remained sufficient for reliable super-resolution imaging, highlighting the exceptional robustness of the self-blinking mechanism and its compatibility with chemically sensitive workflows such as expansion microscopy (ExM) ^16^ .

### Self-blinking nanobody probes are compatible with diverse SMLM modalities

Because different single-molecule localization microscopy (SMLM) modalities impose distinct constraints on fluorophore photophysics, photon budgets, and temporal dynamics, we next evaluated whether self-blinking nanobody probes could serve as a general labeling strategy across complementary localization techniques.

Using the same antibody–secondary nanobody labeling strategy as for wide-field sb-dSTORM, we performed MINFLUX^17^ imaging of microtubules and vimentin filaments labeled with JF635b-conjugated nanobodies. Under identical sample preparation conditions, JF635b-supported MINFLUX localization yielded mean combined localization precisions of 0.90 nm for microtubules and 0.97 nm for vimentin (Fig. 4a, b), demonstrating that the spontaneous blinking kinetics of JF635b are fully compatible with high-precision MINFLUX localization.

**Figure 4.**
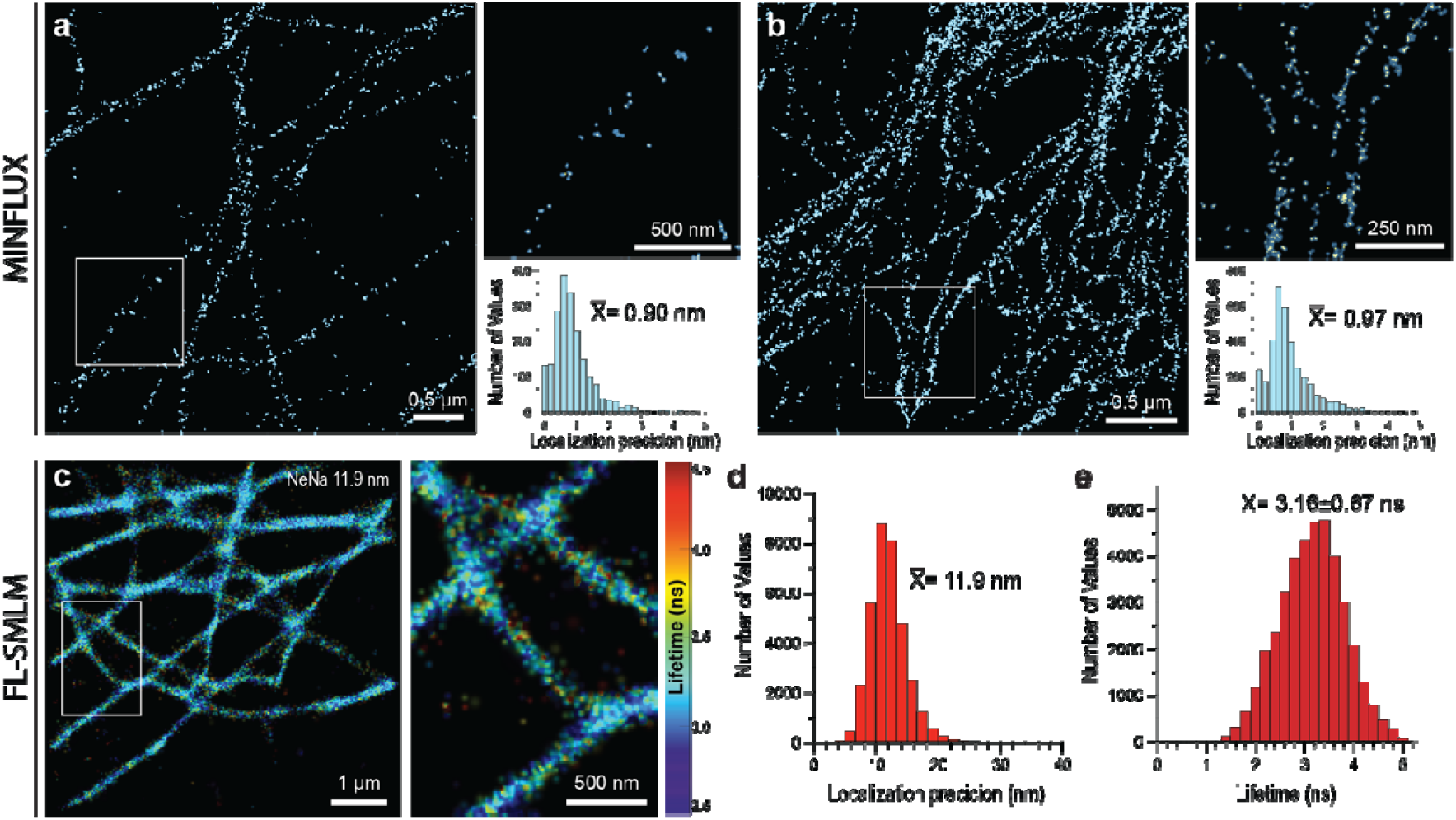
Evaluating JF635b in MINFLUX and fluorescence lifetime SMLM. MINFLUX imaging of the microtubule (**a**) and vimentin (**b**) staining using 2.Nb-JF635b. Combined localization precision plots yield mean precisions of 0.90 nm (microtubules) and 0.97 nm (vimentin). (**c**) Confocal fluorescence lifetime SMLM imaging of microtubules labeled with 1.Ab and 2.Nb-JF635b. (**d**) localization precision histogram, and (**e**) fluorescence lifetime distribution (mean ± s.d. indicated).

We further tested the applicability of self-blinking nanobody probes in fluorescence-lifetime SMLM (FL-SMLM)^12^ using confocal laser scanning microscopy. JF635b-labeled microtubules produced high-quality lifetime-resolved super-resolution reconstructions with a mean localization precision of 11.9 nm and a fluorescence lifetime of 3.16 ± 0.67 ns (Fig. 4c– e). These results demonstrate that JF635b is fully compatible with confocal SMLM while simultaneously providing robust lifetime information, enabling applications such as lifetime-based multiplexing.

Together, these data demonstrate that self-blinking nanobody probes are compatible with a broad range of SMLM implementations, spanning wide-field, confocal, and MINFLUX-based localization microscopy, and can serve as a unified labeling strategy across modalities.

## Discussion

dSTORM relies critically on fluorophore blinking, yet most implementations remain dependent on specialized chemical environments that limit robustness, reproducibility, and long-term sample stability. Although nanobodies offer clear advantages in labeling accuracy and epitope accessibility, reducing linkage error from ∼20-25 nm for conventional antibodies to ∼3 nm, their practical use in SMLM has been constrained by the poor blinking performance of commonly used dyes when directly conjugated. Our results identify fluorophore blinking as a central bottleneck in nanobody-based dSTORM and demonstrate that this limitation can be effectively overcome using self-blinking fluorophores.

By conjugating nanobodies to the spontaneously blinking dye JF635b, we enable robust and sustained localization microscopy under buffer-independent conditions. Self-blinking nanobody probes restore high localization densities and nanometer-scale precision for both direct and secondary labeling strategies, eliminating the need for reducing or oxygen-scavenging buffers. This simplification not only enhances experimental robustness but also lowers technical barriers, particularly in multi-user imaging facilities and high-throughput settings where reproducibility across experiments and users is critical.

A defining feature of JF635b-labeled nanobodies is their exceptional photophysical stability. Stable blinking in pure water and sustained performance after long-term storage for up to one year highlight the intrinsic robustness of the self-blinking mechanism. In contrast to conventional dSTORM approaches, where reactive imaging buffers degrade over time and introduce variability, samples labeled with JF635b can be stored in simple PBS and reimaged over extended periods with minimal loss of performance. This enables repeated imaging of the same specimen under comparable conditions, supporting longitudinal studies, large-scale comparisons, and the development of standardized reference samples. By removing buffer-related variability, self-blinking dSTORM directly addresses a major limitation in SMLM: reproducibility across samples and laboratories.

Beyond long-term stability, JF635b can sustain efficient blinking even in pure water, expanding SMLM’s operational range. Unlike conventional fluorophores that require carefully tuned environments, JF635b blinks reversibly in the absence of salts or additives, enabling imaging with minimal perturbation and simplifying workflows. This also opens opportunities for integration with chemically sensitive techniques such as expansion microscopy, where buffer composition affects structural preservation. Self-blinking nanobody probes work across various SMLM methods, including wide-field self-blinking dSTORM, fluorescence-lifetime SMLM, and MINFLUX, achieving localization precisions below a nanometer. Their cross-modality compatibility makes them a unifying labeling strategy for super-resolution microscopy, providing consistent probe behavior across platforms.

In summary, self-blinking nanobody probes combine minimal linkage error, photophysical robustness, and operational simplicity. By restoring the practical advantages of nanobody labeling and removing buffer dependence, they provide a broadly enabling solution that enhances reproducibility, flexibility, and accessibility in nanoscale super-resolution microscopy.

## Methods

### Cell culture

COS-7 cells were cultured in Petri dishes at 37 °C with 5% CO2 inside a humidified incubator. They were grown in complete Dulbecco’s Modified Eagle Medium (DMEM, ThermoFisher Scientific), supplemented with 10% fetal bovine serum (FBS, ThermoFisher), 4 mM L-glutamine, and 1% penicillin/streptomycin (ThermoFisher). For sample preparation, cells were seeded onto 18 mm poly-L-lysine-coated coverslips in 12-well plates and incubated at 37 °C with 5% CO2 in a humidified environment. For transfection, a plasmid encoding TOM70-EGFP-ALFA-tag driven by a CMV promoter was mixed with Lipofectamine 2000 (Thermo Fisher) and added to the seeded cells according to the manufacturer’s instructions. Transfection and expression were typically left for 16-18h, before cell fixation.

### Immunostaining of cells

COS-7 cells were first treated for 30 s with pre-warmed Extraction Buffer (EB, 10 mM MES, 138 mM KCl,3 mM MgCl_2_, 2 mM EGTA, 320 mM sucrose, pH 6.8) supplemented with 0.1% Saponin. This was directly followed by fixation in a pre-warmed solution of 4% paraformaldehyde and 0.1% Glutaraldehyde in EB at RT for 15 min. After removing the fixatives and rinsing with PBS, unreacted aldehyde groups were quenched using 100 mM glycine in PBS for 15 min. Cells were blocked and permeabilized with 3% serum albumin (BSA) and 0.1% Triton X-100 (Sigma-Aldrich) in PBS for 30 min at RT. For Fig. 1, NbGFP and NbALFA labeled with AF647 or JF635b were purchased as FluoTag^®^-X2 from NanoTag Biotechnologies (clone 1H1, Cat#: N0304; Cat#: N1502) and used at a 1:1000 dilution for 45 minutes. Recombinant anti-ALFA antibody carrying a mouse IgG1 Fc domain (NanoTag Biotechnologies, Cat# N1582) was used at a 1:500 dilution and detected by Goat anti-Mouse IgG (H+L) Cross-Adsorbed Secondary Antibody, Alexa Fluor™ 647, also used in a 1:500 dilution (ThermoFisher, Cat# A-21235). Microtubules and vimentin structures were labeled using a one-step immunofluorescence^15^ with a mouse monoclonal (SySy, Cat# 302211) and a rabbit monoclonal (Abcam, Cat# ab92547), respectively. Secondary detection was performed using FluoTag^®^-X2 anti-Mouse IgG1 (Cat# N2002) or FluoTag^®^-X2 anti-Rabbit IgG (Cat# N2402). See Supp. Table S3 for details on the used Abs and Nbs. All samples, after staining and washing, were post-fixed in 4% PFA for 15-20 minutes at room temperature, the fixative was removed, and the samples were quenched again with 100 mM glycine in PBS.

### Wide-field sb-dSTORM imaging and data analysis

Wide-field sb-dSTORM measurements were performed on a custom-built setup as described previously^5^ and in the Supplementary Information (Supp Figure S1). Regions of interest (ROIs) of up to 50 × 50 µm^2^ were recorded with a pixel size in the object plane of 103.5 nm, an exposure time of 30 ms, and an EMCCD gain of 500. Each dataset consisted of 50,000 frames. The laboratory temperature was maintained at 23.0°C to ensure thermal stability of the setup. Linear sample drifts of typically 1-2 pixels were observed during acquisition and were corrected by a cross-correlation algorithm. Excitation was performed using a highly inclined and laminated optical sheet (HILO) configuration with a laser power of ∼20 mW, yielding an optimal signal-to-noise ratio. Data analysis has been performed using free software FiJi^18^, plugin ThunderSTORM^19^. Same analysis parameters were applied as in previous works^4,5^. Photostability was quantified by tracking the number of localizations per frame over time (Fig. 1d). Localization tables were exported from the ThunderSTORM plugin as .csv files and processed with a custom Python script to generate photostability plots (cumulative localizations in time, see Fig. 1d). The photostability curves were fitted with an exponential decay, from which characteristic photobleaching times were extracted. Segmentation of reconstructed super-resolution images was carried out using a standard Python image pipeline. Images were preprocessed using intensity filtering and morphological operations to ensure filament continuity and remove non-filament objects, and then inverted to enable analysis of voids rather than filaments. Void regions were then identified (and scaled to 10.35 nm/pixel), border-adjacent partial objects were removed, and regions were hole-filled to avoid biasing the measurement of their spacing area. Quantitative parameters, including void area distributions, were extracted (see Supplementary Information (Fig. S3) for details).

### Confocal FL-SMLM sb-dSTORM imaging and data analysis

CLSM measurements were performed using a custom-built confocal microscope as described elsewhere^12,20^ and in the Supplementary Information (Fig. S6). Regions of interest (ROIs) of 7 × 7 µm^2^ were scanned, each subdivided into 70 × 70 virtual pixels (100nm/pixel). Similar image acquisition parameters were used as described previously^12^. During analysis, temporal binning of four consecutive frames was applied to improve signal-to-noise ratio. Data analysis has been performed using custom Matlab-based software TrackNTrace LifetimeEdition^12^, freely available at https://github.com/scstein/TrackNTrace/releases/tag/v2.0.

### MINFLUX sb-dSTORM imaging

MINFLUX sb-dSTORM measurements were performed on a commercial MINFLUX nanoscope (Abberior Instruments GmbH, Göttingen) controlled by *Imspector* software. Data were acquired using the standard 2D acquisition protocol described by Sahl *et al*.^21^, see Supplementary Table S2 for details. Gold colloid particles (150 nm; BBI Solutions, catalogue no. EM.GC150/4, Crumlin, UK) were added to the samples as fiducial markers for active beam-path drift correction.

Regions of interest (ROIs) of 5 × 5 µm^2^ were acquired over a total measurement time of ∼6 h. Excitation was performed with the 642 nm laser at 5-10% output (∼25-50 µW) as part of the MINFLUX localization routine. Image data were saved as MSR files and processed in MATLAB (MathWorks, USA).

Localization precisions were calculated as described previously^22^. In brief, localizations from binding events with more than four detections were combined. For enhanced visualization, MINFLUX images were rendered in Blender 3.4.1: raw localizations were displayed as overlapping blue spheres (20 nm) to illustrate the spread of individual events, while combined localizations were rendered as gold spheres (10 nm) to indicate the most likely positions of fluorophores.

## Data Availability

All data supporting the conclusions of this study are available from the corresponding authors upon reasonable request.

## Code availability

FiJi: https://fiji.sc/

Fiji ThunderSTORM plugin: https://github.com/zitmen/thunderstorm

TrackNTrace https://github.com/NRadmacher/TrackNTrace/tree/ISM;

Abberior Imspector commercially available from Abberior Instruments GmbH, Göttingen, Germany

## Acknowledgements

The authors are grateful to Jannik Hentze for their help with sample preparation and labeling. The authors thank Dr. Peter Ilgen (Fraunhofer ITMP, Göttingen) for MINFLUX data visualization using Blender. The authors are grateful to Dr. Antonio Politi (MPI-NAT, Göttingen) for advice in developing the segmentation analysis pipeline. The authors thank Dr. Steffen Frey (NanoTag Biotechnologies GmbH) for advice. We thank Prof. Philip Tinnefeld’s lab (LMU, Munich) for support with JF635b blinking experiments in water. The authors are grateful to Dr. Ingo Gregor for technical assistance.

## Additional information

Supplementary information is available for this paper.

