## Supplementary material for "Self-blinking dye restores efficient use of nanobodies in single-molecule localization microscopy": Supp. Information

### Wide-field single-molecule localization microscopy

*Optical setup description*

Wide-field measurements were performed using a custom-built optical setup, described elsewhere[1] and shown in Figure S1. A pulsed super-continuum white light laser (WL Laser) (SuperK Fianium, NKT Photonics) was used for sample excitation. A variable filter (VF) (SuperK Varia, NKT Photonics) connected to the laser output enabled flexible selection of the output wavelength (605-652 nm for efficient excitation of JF635b and Alexa Fluor 647). A neutral density filter (NE10A-A, Thorlabs), in tandem with a variable neutral density filter (ND) (NDC-50C-4-A, Thorlabs), was used to adjust the laser excitation power.

The laser beam was coupled into a single-mode optical fiber (SMF) (P1-460B-FC-2, Thorlabs) with a typical coupling efficiency of 40%. After exiting the fiber, the collimated laser beam was expanded 3.6× using telescope lenses (TL1 and TL2). The typical excitation intensity at the sample was approximately 1 $kW/cm^2$, sufficient to switch the fluorophore (Alexa 647) from the bright to the dark state (see zoom-in box in Figure S1) in the presence of the blinking agent cysteamine (30070, Sigma-Aldrich) at 50 mM in PBS. Identical imaging parameters were used for imaging of samples labeled with JF635b.

The laser beam was focused onto the back focal plane of the TIRF objective (UAPON 100X oil, 1.49 NA, Olympus) using an achromatic lens (L1) (AC508-180-AB, Thorlabs). Beam displacement relative to the optical axis for switching between EPI, HILO, and TIRF illumination schemes was achieved using a translation stage (TS) (LNR25/M, Thorlabs). Smooth lateral positioning of the sample was enabled by a high-performance two-axis linear stage (M-406, Newport). In addition, an independent one-dimensional translation stage (LNR25/M, Thorlabs) combined with a differential micrometer screw (DRV3, Thorlabs) was used to move the objective along the optical axis for focusing.

Spectral separation of the collected fluorescence light from the excitation path was achieved using a multi-band dichroic mirror (DM) (Di03 R405/488/532/635, Semrock), which directed the fluorescence light toward the tube lens (L2) (AC254-200-A-ML, Thorlabs). The field of view was physically limited in the emission path by an adjustable slit aperture (SP60, OWIS) positioned in the image plane. Lenses L3 (AC254-100-A, Thorlabs) and L4 (AC508-150-A-ML, Thorlabs) re-imaged the emitted fluorescence light from the slit onto an emCCD camera (iXon Ultra 897, Andor). Band-pass filters (BP) (BrightLine HC 692/40) were used to further block scattered excitation light. The total magnification of the optical system on the emCCD camera was 166.6×, resulting in an effective pixel size of 103.5 nm in the sample space. All experiments were conducted at 23 °C to ensure mechanical stability of the optical setup.

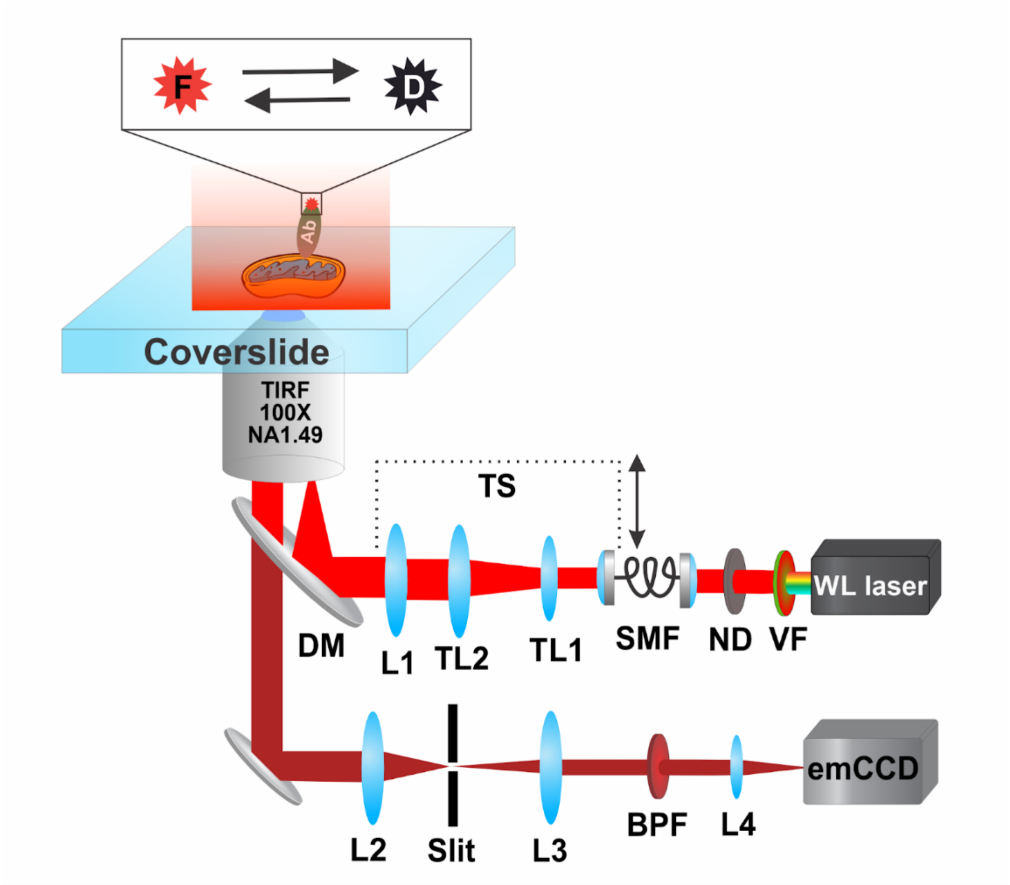


**Figure S1.** Schematics of a custom-built wide-field SMLM setup used for dSTORM imaging. The setup is equipped with a white-light laser source, enabling flexible excitation of fluorophores across different spectral regions, and a scientific emCCD camera with single-molecule sensitivity. It supports imaging using EPI, HILO, and TIRF excitation schemes.

*Experimental conditions and image analysis*

The following imaging parameters were used for both dSTORM and sb-dSTORM: A laser power of 26 mW (measured at the back of the objective) was used to excite both Alexa Fluor 647 (in the presence of GLOX–MEA, 50 mM) and JF 635b (in PBS buffer). Time series of 30,000–50,000 frames were recorded using a gain of 500 and an exposure time of 30 ms. The pixel size in the image plane was 103.5 nm.

The data were analyzed using the ImageJ plugin ThunderSTORM, and the super-resolved images were reconstructed at 10× magnification, yielding a pixel size of 10.35 nm. The reconstructed images were corrected for drift using the cross-correlation algorithm implemented in ThunderSTORM. Finally, filters were applied to reject out-of-focus emitters and poorly fitted Gaussians.

| Target | Antibody/ Nanobody | Dye | Laser [nm] | Laser Power [mW] | Buffer | Total number of Frames | Total imaging time [min] |
|---|---|---|---|---|---|---|---|
| Mitochondria | NbALFA | AF647 | 640 | 35 | GLOX | 50,000 | 25 |
| Mitochondria | NbGFP | AF647 | 640 | 35 | GLOX | 50,000 | 25 |
| Mitochondria | NbALFA | JF635 | 640 | 26 | PBS | 50,000 | 25 |
| Mitochondria | NbGFP | JF635 | 640 | 26 | PBS | 50,000 | 25 |
| Mitochondria | anti-ALFA Ab | AF647 | 640 | 35 | GLOX | 50,000 | 25 |
| Microtubule | αMouse Nb | JF635 | 640 | 26 | PBS | 50,000 | 25 |
| Microtubule | αMouse Nb | JF635 | 640 | 26 | Water | 50,000 | 25 |
| Vimentin | αRabbit Nb | JF635 | 640 | 26 | PBS | 50,000 | 25 |
| PMP70 | αRabbit Nb | JF635 | 640 | 26 | PBS | 50,000 | 25 |

**Supplementary Table S1.** Parameters used for Wide-field data acquisition.

### Microtubules labeled with anti-mouse IgG1-JF635b – Images library

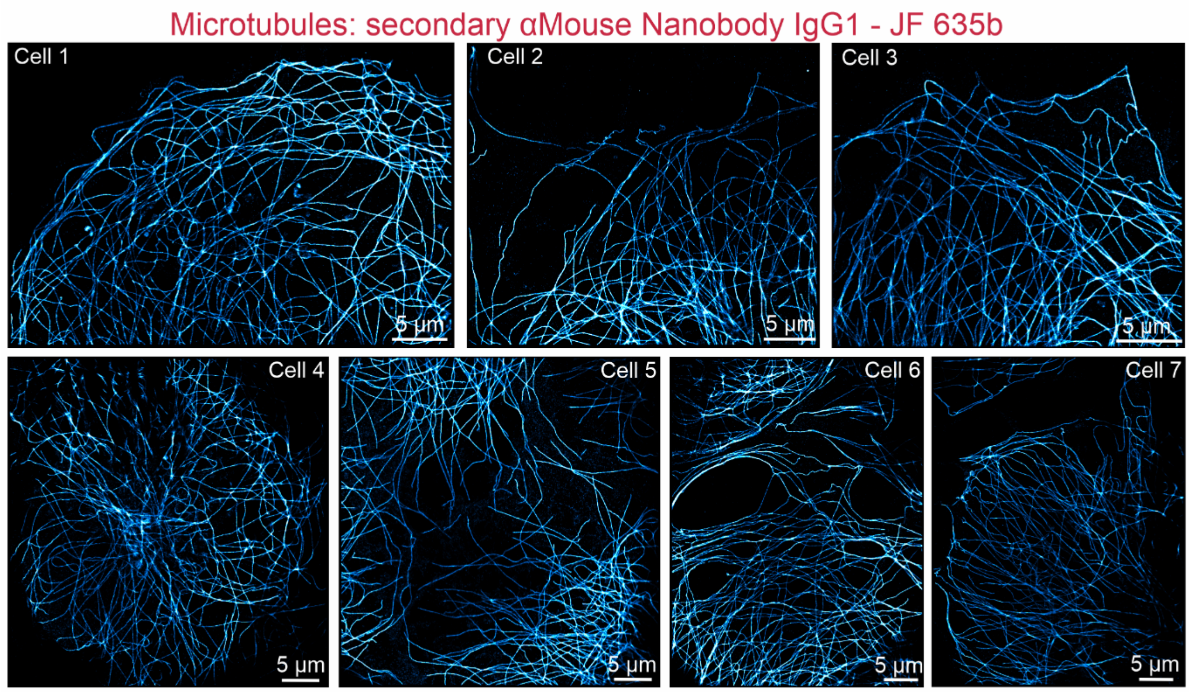


**Figure S2.** Library of sb-dSTORM images of microtubules labeled with JF635b in COS-7 cells imaged in PBS buffer. Representative wide-field sb-dSTORM reconstructions of microtubule networks acquired under identical imaging conditions. Images illustrate the consistency of labeling, filament continuity, and blinking performance across multiple cells and independent preparations.

**Segmentation analysis**

A custom Python-based pipeline was used to segment reconstructed super-resolution images. The workflow consisted of the following steps:

1. **Calibration** – Images were loaded and scaled to the correct dimensions (10.35 nm/pixel).
2. **Pre-filtering** – For microtubule datasets, an initial intensity threshold of 1% was applied.
3. **Inversion** – Images were inverted to emphasize voids rather than filaments.
4. **Morphological opening** – An opening operation was applied using a 3-pixel disk filter.
5. **Void segmentation** – Voids were detected and labeled.
6. **Artifact removal** – Partial voids adjacent to image borders were excluded to avoid bias.
7. **Hole filling** – holes, corresponding to filament fragments, were filled in detected void regions in order not to bias the measurement of mesh spacing.
8. **Quantitative analysis** – Morphometric parameters were extracted, including void area distributions and major/minor axis lengths.

The Python code used for segmentation is available from the corresponding author upon reasonable request.

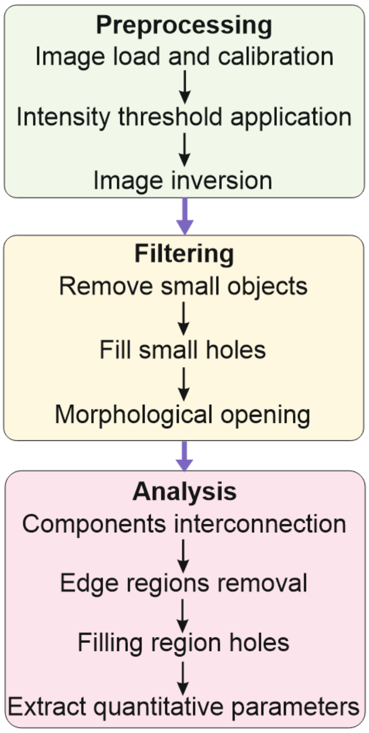


**Figure S3.** Schematics for the segmentation image analysis workflow.

**Vimentin filaments labeled with anti-rabbit nanobody-JF635b – Images library**

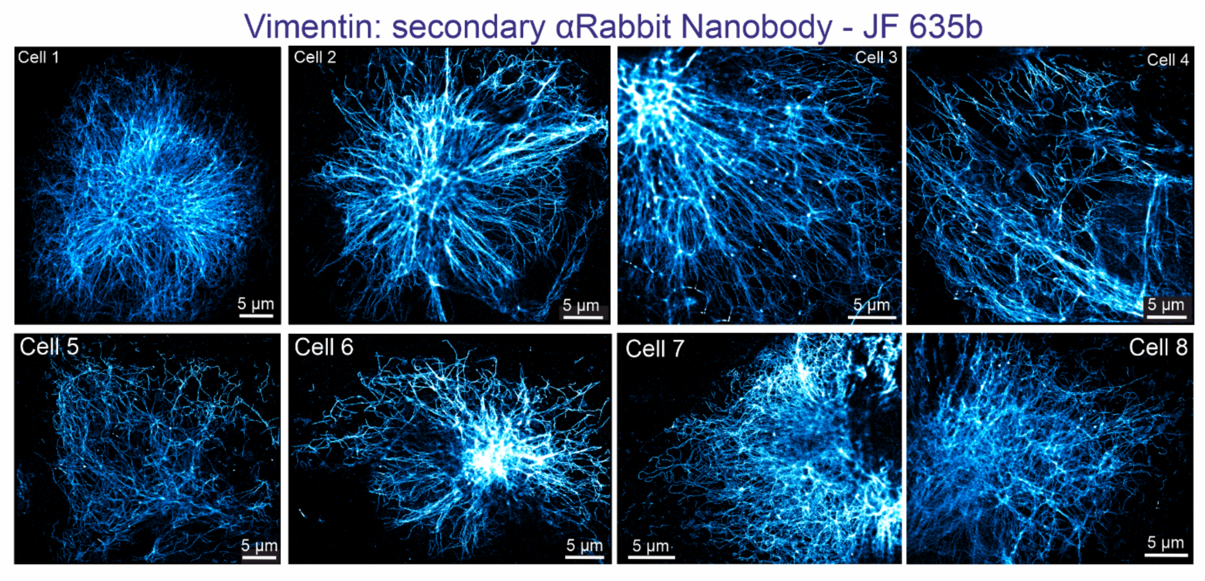


**Figure S4.** Library of sb-dSTORM images of vimentin labeled with JF635b in COS-7 cells imaged in PBS buffer. Representative wide-field sb-dSTORM reconstructions of vimentin intermediate filaments. The images highlight the dense filament networks, uniform labeling, and robust blinking behavior obtained with JF635b-conjugated nanobodies across samples.

**Peroxisomes (PMP70) labeled with anti-rabbit nanobody-JF635b – Images library**

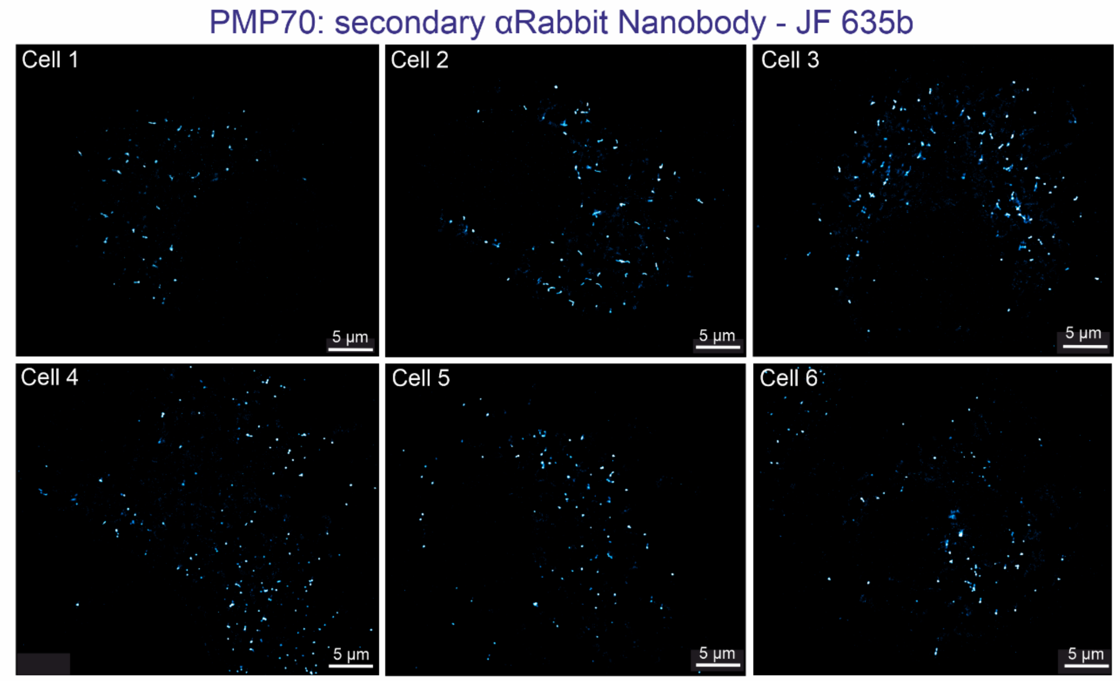


**Figure S5.** Library of sb-dSTORM images of PMP70-labeled peroxisomes in COS-7 cells imaged in PBS buffer. Representative sb-dSTORM reconstructions showing the morphology and distribution of PMP70-positive peroxisomes. The dataset demonstrates that JF635b performs reliably across different cellular targets, labeling strategies, and structural contexts.

### MINLFLUX imaging parameters

| Parameter | MINFLUX iteration | | | | |
|---|---|---|---|---|---|
| | Pre-local. | 1 | 2 | 3 | 4 |
| L (nm) | 289 | 289 | 152 | 76 | 40 |
| TCP | hexagon | hexagon | hexagon | hexagon | hexagon |
| # of photons | 160 | 150 | 100 | 100 | 150 |
| Laser power factor | 1 | 1 | 2 | 4 | 6 |
| TCP dwell time (ms) | ≥1 | ≥1 | ≥1 | ≥1 | ≥1 |
| Pattern repeat factor | 1 | 5 | 5 | 5 | 5 |
| CFR limit | 2,00 | 0,50 | 2,00 | 0,80 | 2,00 |
| Background threshold | 15 | 10 | 10 | 10 | 10 |

**Supplementary Table S2.** Parameters used for MINFLUX data acquisition.

### Confocal single-molecule localization microscopy

*Optical setup description*

CLSM measurements were performed using a custom-built confocal microscopy setup described previously[3]; see Figure S2. Briefly, a white light (WL) laser (Koheras SuperK Power) with a fixed repetition rate of 40 MHz was used. An acousto-optic tunable filter (AOTF) (Koheras SpectraK Dual) attached to the WL laser output provided full flexibility in selecting the excitation wavelength.

The laser beam was coupled into a single-mode fiber (SMF) (PMC-460Si-3.0-NA012 3APC-150-P, Schäfter+Kirchhoff) using a fiber coupler (60SMS-1-4-RGBV-11-47, Schäfter+Kirchhoff), and was decoupled and collimated using a 10× air objective (UPlanSApo 10×, 0.40 NA, Olympus). A quad-band dichroic mirror (DM) (ZT405/488/561/640rpc, Chroma) was used to direct the excitation light into the specimen and to separate it from the emission light. The excitation beam passed through a fast laser scanning system (FLIMbee, PicoQuant GmbH). The scanning system deflected the beam to image large regions of interest while maintaining the focus position by directing the laser beam onto the back focal plane of the objective (UApo N 100×, 1.49 NA oil, Olympus). The region of interest and focus plane were controlled by a manual XY stage (Olympus) and a z-piezo stage (Nano-ZL100, MadCityLabs), respectively. The emitted fluorescence light was collected using the same objective and focused onto a pinhole with a 100 μm diameter (PH) (P100S, Thorlabs) using a 180 mm achromatic lens (L1) (AC508-180-AB, Thorlabs). The emission light was then collected and collimated by a 100 mm lens (L2) (AC508-200-A, Thorlabs). Long-pass filters (LP) (488 LP, 647 LP Edge Basic, Semrock) were used to block excitation laser light in the emission path. Band-pass filters (BP) (BrightLine HC 692/40, Semrock) was used to further reject scattered excitation light.

Finally, the emission light was focused onto a SPAD detector (SPCM-AQRH, Excelitas) using an achromatic lens (L3) (AC254-030-A-ML, Thorlabs). The output signal from the photon detector was recorded using a TCSPC system (HydraHarp 400, PicoQuant).

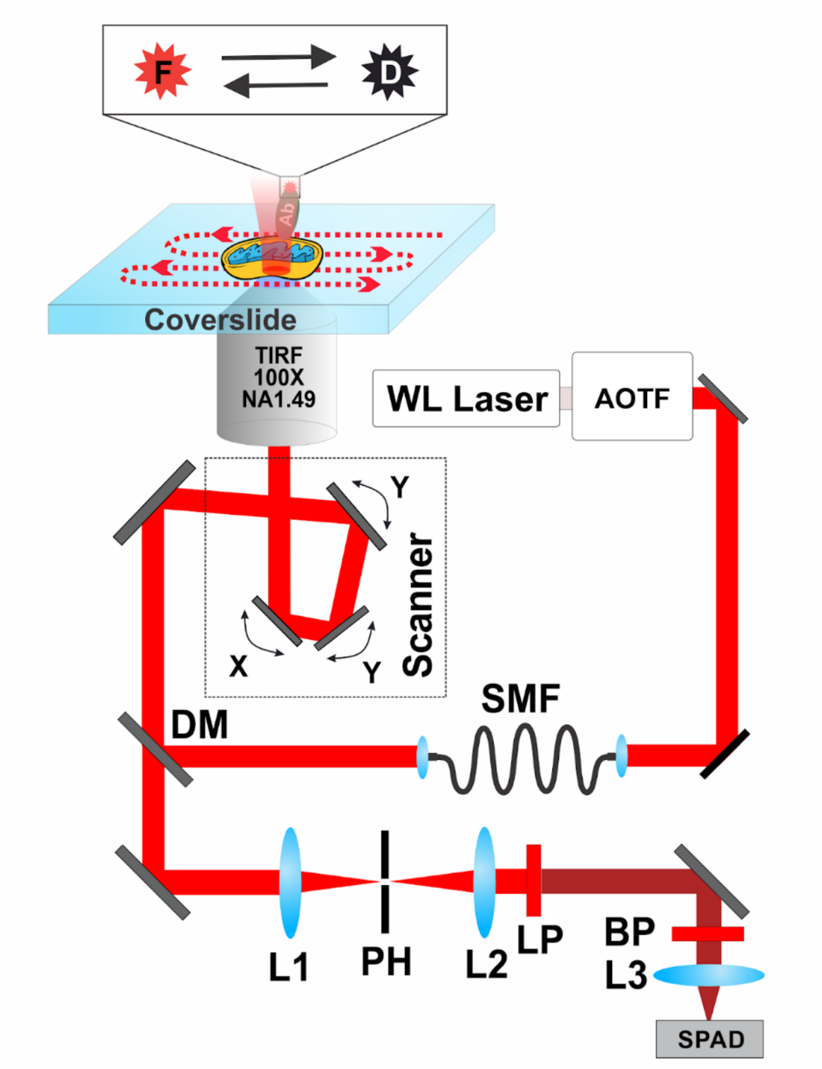


**Figure S6.** Schematic representation of the custom-built time-resolved CLSM optical setup. The setup is equipped with a white light laser source and a time-resolved modality.

*Experimental conditions and image analysis*

Data were acquired using commercial software (PicoQuant SymPhoTime 64), which controlled both the TCSPC and scanner systems. For CLSM-based FL-SMLM, regions of interest measuring 10 × 10 μm² were scanned. Each image consisted of four repeated scans of the region, with a virtual pixel size of 100 nm and a dwell time of 30 μs/pixel. A time series of 50.000 frames was recorded.

| Antibody or Nanobody | Vendor | Information | Dilution |
|---|---|---|---|
| FluoTag®-X2 anti-GFP (NbGFP; clone 1H1) | NanoTag Biotechnologies | Cat#: N0304 | 1:1000 |
| FluoTag®-X2 anti-ALFA-tag (NbALFA) | NanoTag Biotechnologies | Cat#: N1502 | 1:1000 |
| Mouse anti-alpha-Tubulin | Synaptic Systems | Cat# 302211 | 1:1000 |
| Rabbit monoclonal anti-Vimentin | Abcam | Cat# ab92547 | 1:500 |
| Recombinant anti-ALFA Antibody (mouse IgG1) | NanoTag Biotechnologies | Cat# N1582 | 1:500 |
| Goat anti-Mouse IgG Cross-Adsorbed Antibody - AF647 | ThermoFisher | Cat# A-21235 | 1:500 |
| FluoTag®-X2 anti-Mouse IgG1 | NanoTag Biotechnologies | Cat# N2002 | 1:500 |
| FluoTag®-X2 anti-Rabbit IgG | NanoTag Biotechnologies | Cat# N2402 | 1:500 |

**Supplementary Table S3. Antibodies and Nanobodies used for labelling**
